# Germ granules prevent accumulation of somatic transcripts in the adult *C. elegans* germline

**DOI:** 10.1101/089433

**Authors:** Andrew Kekūpa’a Knutson, Thea Egelhofer, Andreas Rechtsteiner, Susan Strome

## Abstract

The germ cells of multicellular organisms protect their developmental potential through specialized mechanisms. A shared feature of germ cells from worms to humans is the presence of non-membrane-bound ribonucleoprotein organelles called germ granules. Depletion of germ granules in *Caenorhabditis elegans* (i.e., P granules) leads to sterility and in some germlines expression of the neuronal transgene *unc-119::gfp* and the muscle myosin MYO-3. Thus, P granules are hypothesized to maintain germ cell totipotency by preventing somatic development, although the mechanism by which P granules carry out this function is unknown. In this study, we performed transcriptome and single molecule RNA-FISH analyses of dissected P-granule-depleted gonads at different developmental stages. Our results demonstrate that P granules are necessary for adult germ cells to down-regulate spermatogenesis RNAs and to prevent the accumulation of numerous soma-specific RNAs. P-granule-depleted gonads that express the *unc-119::gfp* transgene also express many other genes involved in neuronal development and concomitantly lose expression of germ cell fate markers. Finally, we show that removal of either of two critical P-granule components, PGL-1 or GLH-1, is sufficient to cause germ cells to express UNC-119::GFP and MYO-3 and to display RNA accumulation defects similar to those observed after depletion of P granules. Our data identify P granules as critical modulators of the germline transcriptome and guardians of germ cell fate.

## INTRODUCTION

A cell’s fate and function are determined by its transcriptome and proteome. The transcriptomes of germ cells are unique and are comprised of RNAs that encode proteins essential for specialized, germ cell-specific functions, such as meiosis, spermatogenesis, and oogenesis (Lesch and Page 2012). Throughout development, germ cells must maintain expression of germline-specific genes as well as repress genes that would lead to somatic differentiation (Strome and Updike 2015). Failure to maintain appropriate gene expression programs can lead to sterility, preventing an organism from passing its genetic information to future generations.

A conserved feature of germ cells is the presence of electron-dense, non-membrane-bound organelles called germ granules, which include polar granules in *Drosophila*, chromatoid bodies in mouse spermatids, Balbiani bodies in mouse oocytes, and P granules in *C. elegans* (reviewed in Voronina *et al*. 2011). Germ granules are typically associated with the cytoplasmic face of germ cell nuclei and have been observed to overlie clusters of nuclear pores in worms, zebrafish, and mice (Pitt *et al*. 2000; Knaut *et al.* 2000; Chuma *et al*. 2009). Many essential germ-granule factors are known or predicted to bind RNA. Thus, germ granules are suspected to serve as posttranscriptional processing centers to modify, degrade, and/or store mRNAs as they exit germ cell nuclei (Voronina *et al*. 2011; Updike and Strome 2010).

In *C. elegans*, P granules are maternally loaded into the 1-cell embryo and then asymmetrically segregated to the germ lineage during embryogenesis (Updike and Strome 2010). Although segregation of the bulk of maternal P granules to the germline blastomeres is not required for germ cell specification (Gallo *et al.* 2010), P granules are required for proper germ cell development and may protect germ cells from stressful conditions such as high temperature (Kawasaki *et al*. 1998; Kawasaki *et al*. 2004; Spike *et al*. 2008; Gallo *et al.* 2010). The PGL and GLH proteins make up the core constitutive components of P granules. The three PGL proteins are worm-specific. The two most important PGL proteins, PGL-1 and PGL-3, contain a predicted RNA-binding motif at their C-terminus called an RGG (Arg-Gly-Gly) box (Kawasaki *et al*. 1998; Kawasaki *et al*. 2004). The four GLH proteins are homologs of the highly conserved Vasa protein and contain DEAD-box helicase domains, which may also bind and modulate RNAs (Gruidl *et al.* 1996; Kuznicki *et al.* 2000; Spike *et al.* 2008). The most important GLH protein, GLH-1, is necessary for P-granule retention at the nuclear periphery (Updike *et al*. 2011). In addition to the PGL and GLH core components, P granules contain proteins involved in small RNA biogenesis and the RNAi pathway, suggesting that P granules are involved in small-RNA regulation in germ cells (Updike and Strome 2010; Kasper *et al.* 2014).

Recently P granules have been implicated in maintaining germ cell fate (Updike *et al*. 2014). Germ cells that are depleted of the most important P-granule components (PGL-1, PGL-3, GLH-1, and GLH-4) sometimes express a neuron-specific transgene and a muscle-specific myosin, supporting the hypothesis that P granules prevent somatic development in the *C. elegans* germline. These findings raise many questions, including: 1) What is the extent of somatic development in P-granule-depleted germlines? 2) Is any particular somatic fate favored over others? 3) How early do P-granule-depleted germ cells start expressing somatic markers? 4) What is the mechanism by which P granules prevent expression of somatic genes in the germline?

In this study, we investigated the role of P granules in maintaining appropriate transcript accumulation patterns in the *C. elegans* germline. Using transcript profiling and single molecule RNA-FISH of dissected gonads, we show that upon depletion of P granules, major changes in mRNA levels occur in the germline. Those changes include persistence of sperm transcripts past the normal sperm-production period in hermaphrodites and progressive up-regulation of transcripts from numerous genes normally expressed only in somatic cells. P-granule-depleted germ cells express numerous genes involved in neuronal fate and differentiation, and those cells lose their germ cell identity, highlighting the antagonistic relationship between germ and somatic fates. Finally, we show that germlines that lack either PGL-1 or GLH-1 also misexpress somatic markers, suggesting that these factors are critical for P-granule function. Our studies show that a key role of P granules is to maintain germ cell identity, perhaps by regulating the stability and/or transcription of target mRNAs.

## MATERIALS AND METHODS

### Strains and worm husbandry

*C. elegans* strains were maintained at 15° or 20° on OP50 bacteria as described (Brenner 1974). Strains used in this study include SS1174 wild type, DP132 *edIs6[unc119p::unc119::gfp + rol-6] IV*, SS0002 *pgl-1(ct131) him-3(e1147) IV,* SS0804 *glh-1(gk100) I/hT2g(I;III)*, SS1193 *glh-1(ok439) I/hT2g(I;III); edIs6[unc-119p::unc-119::gfp + rol-6] IV*, and SS1162 *glh-1(ok439) glh-4(gk225) I/hT2g(I; III); edIs6[unc-119p::unc-119::gfp + rol-6] IV*.

### Gonad isolation and RNA extraction

For experiments using RNAi, synchronized *unc-119::gfp* L1 worms (P0) were placed on HT115 bacteria containing empty RNAi vector (control) or expressing double strand RNA that targets P granules (i.e., *pgl-1*, *pgl-3*, *glh-1*, and *glh-4*) or targets *pgl-1* or *pgl-1*; *pgl-3* (Updike *et al*. 2014; Campbell and Updike 2015) at 24° (Day 0). Three days later, F_1_ larvae were transferred to new control or P-granule RNAi plates and returned to 24°. The majority of these F_1_ larvae were at the L1 or L2 stage at the time of transfer. The next day (Day 4), L4 worms were visually identified by their size and the presence of a white crescent at their vulva. Gonads were dissected from L4 control worms and L4 P-granule RNAi worms and cut at the gonad bend with 30½ gauge needles in egg buffer (27.5 mM HEPES, 130 mM NaCl, 2.2 mM MgCl_2_, 2.2 mM CaCl_2_, and 0.528 mM KCl) containing 0.5% Tween-20 and 1 mM levamisole. Sterility could not be assessed in L4 worms. However, greater than 80% sterility was observed in P-granule RNAi adults that were aged from the population of worms from which L4 gonads were collected. For Day 1 adults, worms were allowed to age one day past the L4 stage and then dissected. For Day 2 adults, worms were allowed to age two days past the L4 stage and then dissected. In Day 2 adults, gonads that visually expressed the *unc-119::gfp* transgene were collected as a separate sample from those that did not detectably express *unc-119::gfp*. For adult control gonads, gonads were dissected from fully fertile worms and cut at the gonad bend. For adult P-granule RNAi worms, gonads were collected only from sterile worms, as judged by the lack of embryos in the uterus. Large P-granule RNAi gonads were cut at the gonad bend; stunted P-granule RNAi gonads were collected in their entirety. Once dissected, gonad arms were collected in Trizol, and total RNA was extracted. For control samples, 75 to 100 gonads were collected per replicate. For P-granule RNAi samples, 125 to 150 gonads were collected per replicate. Three biological replicates were performed for each treatment and time point.

For experiments using P-granule mutants, wild-type (control), *pgl-1* single mutants, or *glh-1* single mutant adults (isolated from *glh-1(gk100)/hT2g* balanced parents) were allowed to lay eggs overnight on OP50 bacteria at 24°. The day after the egg lay, the adults were removed and the F_1_ progeny were returned to 24° for two days. Day 1 adult gonads were dissected, collected into Trizol, and total RNA was extracted. Only distal gonad arms were collected from fertile wild-type worms or sterile P-granule mutant worms, as judged by the lack of embryos in the uterus. Three biological replicates were performed for the wild-type and *pgl-1* samples, and two biological replicates were performed for the *glh-1* sample.

### Transcript profiling and analysis

Ribosomal RNA was depleted from each RNA sample using an NEBNext rRNA Depletion Kit (Human/Mouse/Rat) (Cat. #E6310), and libraries were constructed using an NEBNext Ultra RNA Library Prep Kit for Illumina sequencing (Cat. #E7530). Libraries were sequenced at the Vincent J. Coates Genomics Sequencing Laboratory at UC Berkeley using the Illumina HiSeq 2000, 2500, and 4000 platforms. Raw sequences were mapped to transcriptome version WS220 using Tophat2 (Kim *et al.* 2013). Only reads with one unique mapping were allowed; otherwise default options were used. Reads mapping to ribosomal RNAs were removed from further analysis. DESeq2 (Love *et al.* 2014) was used for differential expression analysis. A Benjamini-Hochberg multiple hypothesis corrected p-value cutoff of 0.05 was used as a significance cutoff. Read counts for each gene were normalized using the size factor normalization function in DESeq2 to ~3.5 million reads genome-wide per condition, after which replicate values were averaged. Gene Ontology analysis was done using the DAVID Bioinformatics Resource version 6.7.

### Gene categories

Gene categories were defined using published microarray and SAGE datasets that profiled specific tissues or whole worms that contained or lacked a germline (Baugh *et al.* 2003; Reinke *et al.* 2004; Meissner *et al.* 2009; Wang *et al.* 2009), as described in (Rechtsteiner *et al*. 2010; Knutson *et al*. 2016). Ubiquitous genes (1,895 genes) are genes that are expressed (tag > 0) in all SAGE datasets that profiled germline, muscle, neuronal, and gut tissue, but that are not in the germline-enriched category. Germline-enriched genes (2,230 genes) are genes with transcripts enriched in adults with a germline compared with adults lacking a germline as assessed by microarray analysis. Germline-specific genes (169 genes) are genes whose transcripts are expressed exclusively in the adult germline and accumulate in embryos strictly by maternal contribution. Sperm-specific genes (858 genes) are genes in the spermatogenesis-enriched category as described in (Reinke et al. 2004) minus genes in the germline-enriched and germline-specific categories. Soma-specific genes (1,181 genes) are genes expressed (tag > 4) in muscle, neuronal, and/or gut tissue, but not expressed (tag = 0) in germline SAGE datasets and also not in the germline-enriched category.

### Unsupervised clustering analysis

The k-means clustering algorithm in R was used to cluster the log_2_ fold changes shown in Figure S3. 14,571 genes significant in at least one of ten comparisons (shown as fold change comparisons in Figure S3) at an adjusted p-value of 0.05 and a log_2_ fold change of at least one were clustered. The number of clusters was increased until the number of clusters and their patterns stabilized but were distinct, resulting in eight clusters. Expression values in the various conditions are shown as DESeq2 normalized log_10_ read counts; they were scaled by a factor of two, so that they would have a similar range as the fold change values. The DESeq2 size factor normalization scaled the samples to a total of ~3.5 million genome-wide reads per condition. Within each cluster the genes were sorted from highest to lowest average expression values across conditions, but expression values were not used to cluster the genes. Our gene set categories are also shown, red indicating the gene being part of the gene set. Boxplots of DESeq normalized log_10_ read counts for each gene cluster are shown across conditions in Figure S4.

### Single molecule RNA fluorescence in situ hybridization (smFISH)

smFISH analysis was performed on L4, Day 1 Adult, and Day 2 Adult control and P-granule RNAi gonads dissected from *unc-119::gfp* worms as described (Lee *et al*. 2016). Stellaris probes were designed, synthesized, and labeled by Biosearch Technologies. The *let-2* and *aqp-2* probes were coupled to Cal Fluor Red 610, and the *cebp-1*, *puf-9*, and *lag-1* probes were coupled to TAMRA. Each probe was resuspended in 250 μL of TE buffer pH 8.0 and then further diluted for hybridization (*let-2* and *lag-1* probes were diluted 1:10, *aqp-2* 1:30, and *cebp-1* and *puf-9* 1:50). Gonads were imaged on a Solamere spinning disk confocal system controlled by Micro-Manager software (Edelstein *et al.* 2014). The set-up was as follows: Yokogawa CSUX-1 scan head, Nikon TE2000-E inverted stand, Hamamatsu ImageEM x2 camera, 561 nm laser, and Plan Apo 60x/1.4 n.a. oil objective. Figures 4, 6E, S5, and S6B contain montages generated by splicing together contiguous images acquired with identical settings. Control and P-granule RNAi pairs used identical confocal settings. All images were processed with Image J and Adobe Illustrator.

### Immunocytochemistry

For antibody staining, gonads were dissected using 30½ gauge needles in egg buffer containing 0.5% Tween-20 and 1 mM levamisole. Dissected gonads were permeabilized using the freeze-crack method, fixed for 10 min in methanol and 10 min in acetone, and stained as described (Strome and Wood 1983). Antibody dilutions were 1:4000 rabbit anti-UNC-64 (Saifee *et al*. 1998), 1:1000 rabbit anti-REC-8 (Liu *et al.* 2011), 1:500 guinea pig anti-HTP-3 (MacQueen *et al*. 2005), 1:2000 mouse anti-GFP (Roche, Cat. #11814460001), 1:2000 rabbit anti-GFP (Novus, Cat. #NB600-308), 1:2000 mouse anti-MYO-3 (Priess and Thomson 1987), and 1:500 Alexa Fluor secondary antibodies (Life Technologies). Images were acquired on the Solamere spinning disk confocal system described above. Figures 5C, 6A, 6C, and S7B contain montages generated by splicing together contiguous images acquired using identical settings. All images were processed with ImageJ and Adobe Illustrator.

For quantification of REC-8 or HTP-3 signal relative to UNC-119::GFP signal, three optical slices were projected for both control and P-granule RNAi gonads. Using the color histogram tool in ImageJ, the intensity of the red channel (REC-8 or HTP-3 staining) was recorded and plotted as a function of the intensity of the green channel (UNC-119::GFP) from random individual cells within the gonad. For each experiment, one control gonad and three P-granule RNAi gonads were sampled.

### Data availability

Strains and reagents are available upon request. File S1 contains processed gene expression data including fold changes and average read counts. Gene expression data are available at GEO with the accession number XXX.

## RESULTS

### P-granule RNAi effectively depletes P-granule mRNAs from the germline

Depletion of P granules from the *C. elegans* germline causes sterility and, in some sterile worms, expression of the pan-neuronal *unc-119::gfp* transgene (Updike *et al.* 2014). Among UNC-119::GFP(+) gonads, approximately 60% also co-expressed the muscle myosin MYO-3 (Updike *et al.* 2014). UNC-119::GFP and MYO-3 were often observed in the same regions of the germline, and those regions showed morphological changes not typical of germ cells (i.e., loss of the rachis, loss of proper germ cell organization, and abnormal nuclear size). These features raise the possibility that depletion of P granules causes germ cells to undergo large-scale developmental changes in gene expression.

To identify genes that become misregulated upon depletion of P granules and to investigate the extent of reprogramming toward somatic cell types, we performed transcriptome analysis of dissected gonads from P-granule-depleted worms compared to control worms. We treated L1 larvae that harbored the *unc-119::gfp* transgene with P-granule RNAi and manually dissected and collected gonads from their F_1_ progeny at different developmental stages: L4 larvae, Day 1 sterile adults, and Day 2 sterile adults (Figure 1A, see Materials and Methods). Throughout this report, we use the shorthand “L4” to mean the L4 larval stage and “PG(-)” to mean P-granule depleted. Of the Day 2 sterile adults, approximately 10% expressed the *unc-119::gfp* transgene in one or both gonad arms. These “transformed” gonads (i.e., Day 2 Adult PG(-) UNC-119::GFP(+) gonads) were collected as a separate sample from Day 2 gonads that did not detectably express the *unc-119::gfp* transgene (i.e., Day 2 Adult PG(-) UNC-119::GFP(-) gonads; Figure 1B). For comparison, worms were fed empty vector RNAi instead of P-granule RNAi, and control gonads were collected from fertile F_1_ worms at the same developmental stages (L4, Day 1 adult, and Day 2 adult). From control worms, only the distal halves of gonads were profiled by cutting at the gonad bend so as to avoid oocytes, which are not present in P-granule-depleted gonads. From P-granule-depleted worms, large gonads were cut at the gonad bend and stunted gonads were collected in their entirety. Three biological replicates were collected for each treatment at each time point. Total RNA was isolated from these replicates, depleted of ribosomal RNA, and used to construct libraries for next-generation sequencing.

**Figure 1.**
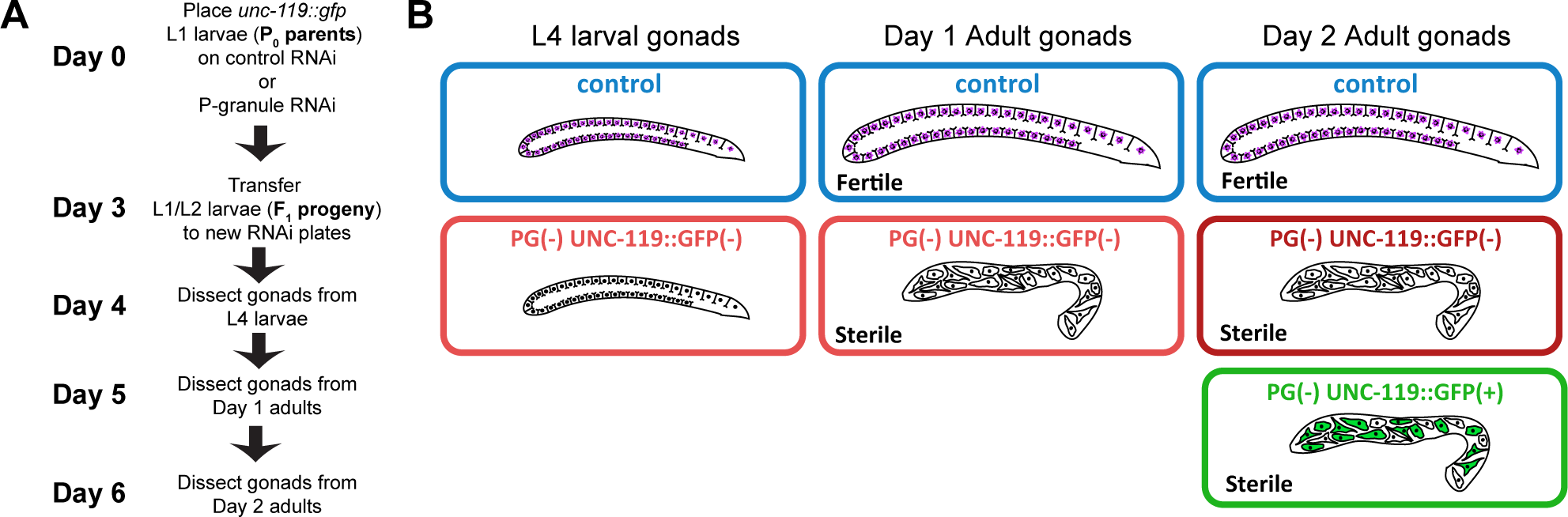
Method to profile transcripts from P-granule-depleted gonads. **A)** Experimental scheme and timeline to analyze the transcriptomes of control and P-granule-depleted gonads from L4 larval, Day 1 adult, and Day 2 adult stages of worms grown at 24^o^. **B)** Seven different samples were collected (three control, and four P-granule RNAi) all in triplicate. PG(-) is shorthand for P-granule depletion. Throughout this report, the samples are color-coded: control samples are in light blue; L4 and Day 1 Adult PG(-) UNC-119::GFP(-) samples are in pink; Day 2 Adult PG(-) UNC-119::GFP(-) samples are in red; and Day 2 Adult PG(-) UNC-119::GFP(+) samples are in green.

Our P-granule depletion method was highly effective. Compared to control gonads, P-granule-depleted gonads at all three developmental stages showed between a 9‐ and 133-fold reduction in *pgl-1*, *pgl-3*, *glh-1*, and *glh-4* transcripts (Figure S1A and B). In addition to the four P-granule targets included in our RNAi construct, our P-granule RNAi also depleted *glh-2* and the DEAD-box helicase domain-containing pseudogene *T08D2.3*, likely due to their extensive sequence identity with *glh-1* and *glh-4*. UCSC genome browser tracks of these P-granule loci revealed read pile-ups spanning approximately 350 base pairs (Figure S1A). Based on their mapped locations, we infer that these reads originate from the RNAi construct. We also observed a 4-fold increase of a portion of *unc-119* transcripts in Day 2 Adult PG(−) UNC-119::GFP(+) gonads compared to Day 2 Adult control gonads (Figure S1C). The reads that mapped to *unc-119* are most likely from the *unc-119::gfp* transgene and not the endogenous *unc-119* locus, since only exons 2, 3, and half of 4 were up-regulated and only those exons are present in the transgene. Taken together, these data show that our P-granule RNAi treatment is highly effective at depleting P-granule mRNAs and that Day 2 Adult PG(-) UNC-119::GFP(+) gonads up-regulate transcription of the *unc-119::gfp* transgene.

### P-granule-depleted adult gonads display progressive changes in transcript accumulation

We observed genome-wide changes in transcript accumulation in P-granule-depleted adult gonads compared to control gonads. Using cut-offs of an adjusted p-value of less than 0.05 and a fold change of more than 2-fold, we identified 2,383 genes as significantly up-regulated and 127 genes as significantly down-regulated in Day 1 Adult PG(-) UNC-119::GFP(-) gonads compared to Day 1 Adult control gonads (Figure 2A). Allowing worms to age to Day 2 of adulthood resulted in 6,833 genes significantly up-regulated and 965 genes significantly down-regulated in Day 2 Adult PG(-) UNC-119::GFP(-) gonads compared to Day 2 Adult control gonads (Figure 2A). Day 2 Adult PG(-) UNC-119::GFP(+) gonads showed even greater transcript accumulation changes when compared to Day 2 Adult control gonads: 8,777 genes significantly up-regulated and 3,049 genes significantly down-regulated (Figure 2A). Comparison of the different up‐ and down-regulated genes from each adult sample revealed high overlap but also unique genes within later gonad samples (Figure 2B). The nested Venn diagrams suggest that gene misexpression in P-granule-depleted adult gonads becomes progressively more severe over time and especially in gonads that express a somatic transgene (Figure 2B).

**Figure 2.**
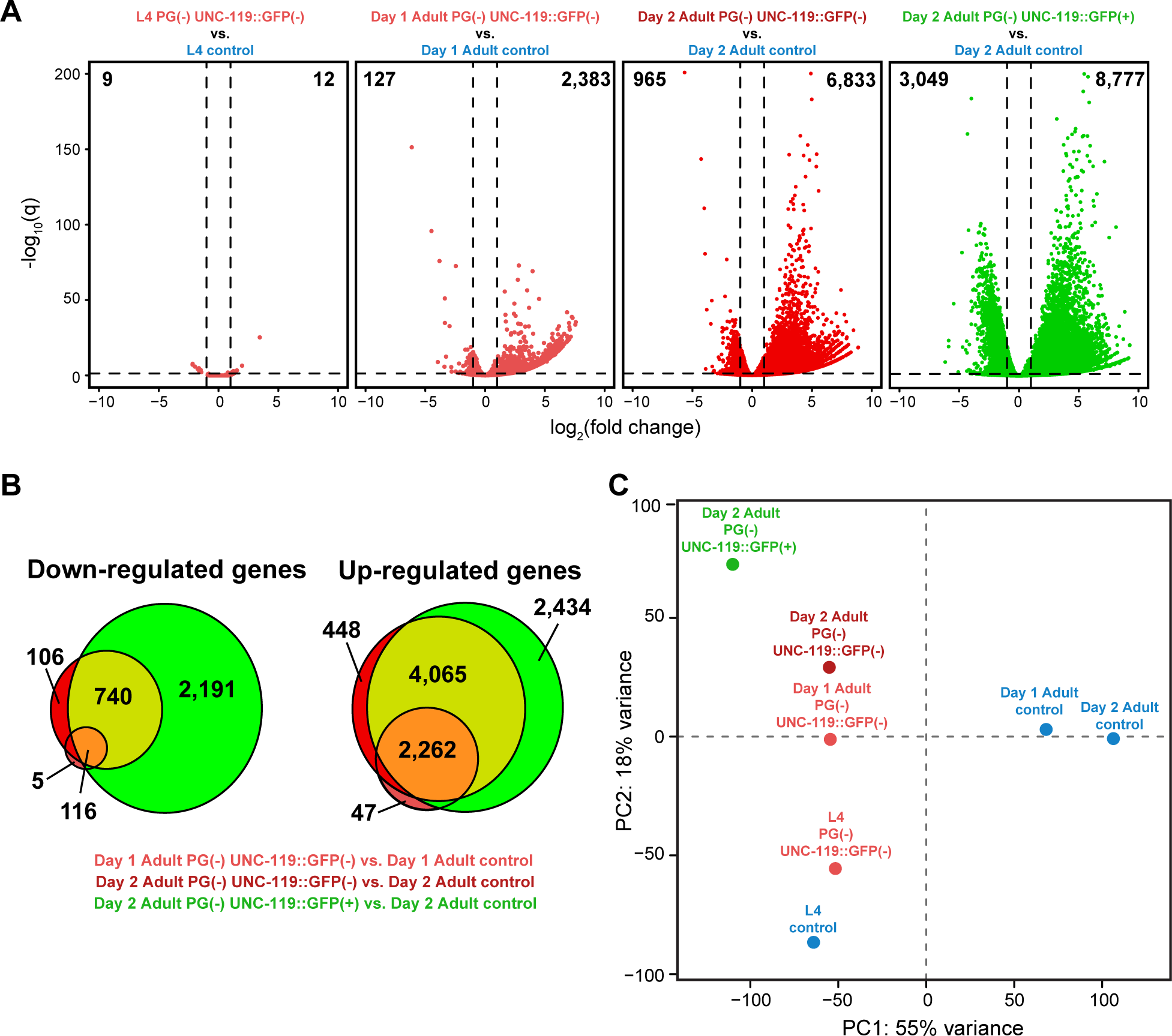
Large, progressive changes in transcript accumulation are observed in adult P-granule-depleted gonads but not in L4 larval gonads. **A)** Volcano plots comparing P-granule-depleted gonads to control gonads at different time points. Dashed lines indicate the significance cut-off of q=0.05 (horizontal line) and a 2-fold change in transcript level (vertical lines). The numbers of significantly misregulated genes in each quadrant are shown at the top of each plot. **B)** Venn diagrams showing the overlap of genes down‐ or up-regulated in Day 1 Adult PG(-) UNC-119::GFP(-) gonads (pink), Day 2 Adult PG(-) UNC-119::GFP(-) gonads (red), and Day 2 Adult PG(-) UNC-119::GFP(+) gonads (green). The overlap of all three samples is shaded in orange. **C)** Principle component analysis (PCA) of all seven samples (P-granule-depleted and control samples) across two principle components (PC1 and PC2).

### P-granule-depleted L4 larval gonads display little change in transcript accumulation

In contrast to Day 1 and Day 2 Adult PG(-) gonads, L4 PG(-) gonads showed essentially no transcript accumulation changes compared to L4 control gonads; only nine genes were significantly down-regulated and 12 genes significantly up-regulated (Figure 2A). Of the nine down-regulated genes, five genes were direct P-granule RNAi targets (i.e., *pgl-1*, *pgl-3*, *glh-1*, *glh-4*, *glh-2*) and two genes were likely the result of 3’ to 5’ transitive RNAi spreading (*him-3* and *hil-4*, which are at the 5’ ends of *pgl-1* and *pgl-3*, respectively; Alder *et al.* 2003). Another gene, *T12F5.2*, is down-regulated as a result of being in the same operon as *glh-4.* The 12 genes significantly up-regulated in P-granule-depleted L4 gonads did not fall into any specific Gene Ontology category.

Principle component analysis (PCA) of all seven samples revealed clustering of the Day 1 and Day 2 Adult control gonad samples close to each other and far from the L4 control sample and all of the PG(-) samples along PC1 (Figure 2C). Surprisingly, the control L4 gonad sample is similar to all of the PG(-) samples along PC1 (Figure 2C). This along with results discussed below suggests that gonads isolated from P-granule-depleted adult worms are developmentally more similar to L4 gonads than adult gonads and possibly maintain mRNAs that are normally restricted to the L4 stage. These results suggest that P granules are necessary for the normal progression of germline development from the L4 stage to adult. The PG(-) samples spread out along PC2, with the Day 2 Adult PG(-) UNC-119::GFP(+) gonad sample being farthest from the others. This likely reflects differences in transcript accumulation that may underlie somatic development in transformed Day 2 Adult PG(-) UNC-119::GFP(+) gonads.

### P-granule-depleted adult gonads down-regulate germline transcripts and up-regulate somatic and sperm transcripts

To determine the types of genes that are misregulated in response to P-granule depletion, we analyzed the levels of mapped reads across different gene categories: ubiquitously expressed, enriched expression in the germline, expressed specifically in the germline, expressed specifically in sperm, or expressed specifically in somatic cells. These categories were defined previously using microarray and serial analysis of gene expression (SAGE) data from different tissues and from animals that possessed or lacked a germline (see Materials and Methods). After normalizing for sequencing depth across all seven samples, we compared average read counts across different gene categories.

Ubiquitous genes displayed constant levels of read counts across all seven samples (Figure 3A). Closer examination of individual genes revealed a small subset of ubiquitous genes that showed up-regulation in P-granule-depleted gonads, especially in Day 2 Adult PG(-) UNC-119::GFP(+) gonads (Figure S2A). However, for the majority of ubiquitous genes, including *ama-1* which encodes the large subunit of RNA Polymerase II, read levels were not changed by P-granule depletion (Figure 3B).

**Figure 3.**
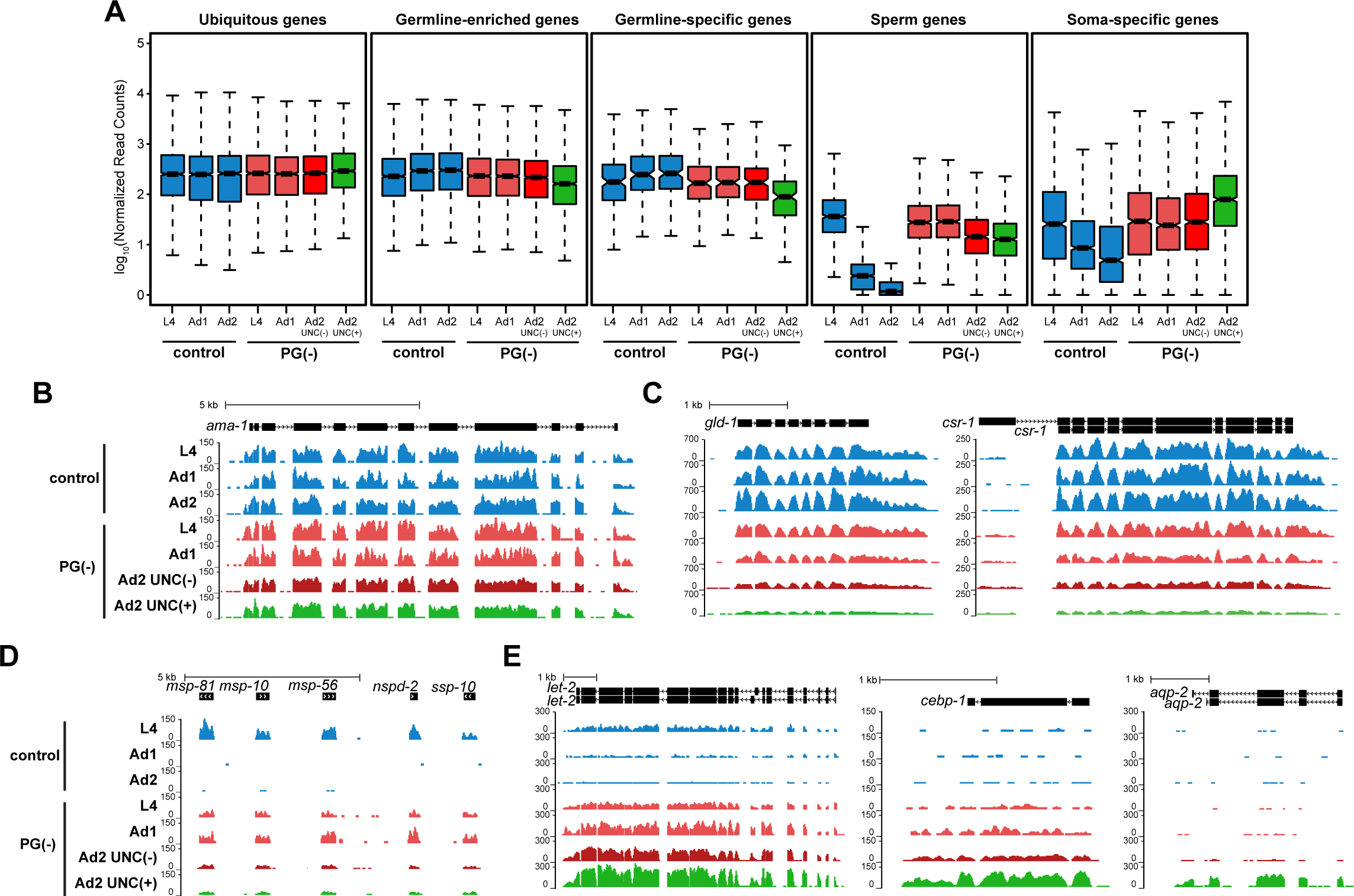
P-granule-depleted gonads down-regulate germline genes and up-regulate sperm and somatic genes. **A)** Boxplots of normalized RNA-seq reads from ubiquitous genes, genes with germline-enriched expression, genes with germline-specific expression, sperm genes, and soma-specific genes in control and P-granule-depleted gonads at different stages. Each box extends from the 25^th^ to the 75^th^ percentile of read values. The whiskers extending from each box indicate the 2.5^th^ and the 97.5^th^ percentiles. Wedges around the median indicate 95% confidence intervals for the medians. **B-E)** UCSC genome browser tracks showing the abundance of RNA-seq reads from control and P-granule-depleted gonad samples across (B) the ubiquitous gene *ama-1*, (C) germline-specific genes *gld-1* and *csr-1*, (D) sperm-specific genes *msp-81*, *-10*, and *-56*, *nspd-2*, and *ssp-10*, and (E) soma-specific genes *let-2*, *cebp-1*, and *aqp-2*. The sample names in the boxplots and the genome browser tracks are abbreviated with L4 denoting L4 larvae, Ad1 denoting Day 1 Adults, and Ad2 denoting Day 2 Adults. Additionally, UNC(-) denotes no expression and UNC(+) denotes expression of the *unc-119::gfp* transgene. The boxplots and browser tracks are color-coded as described for Figure 1.

Germline-enriched and germline-specific gene read counts in control gonads increased progressively from L4 larvae to Day 1 adults to Day 2 adults (Figure 3A). Upon P-granule depletion, L4 PG(-), Day 1 Adult PG(-), and Day 2 Adult PG(-) UNC-119::GFP(-) gonads showed levels of reads across germline genes that were similar to levels observed in L4 control gonads (Figure 3A). Those levels dropped in Day 2 Adult PG(-) UNC-119::GFP(+) gonads compared to Day 2 Adult control gonads (Figures 3A and S2B). Examples of germline transcripts that were down-regulated include transcripts encoding the P-granule-associated argonaute CSR-1 and the P-granule-associated translational repressor GLD-1 (Figure 3C). These data suggest that P granules are required for the normal accumulation of germline transcripts and prevent the decrease in germline mRNAs in gonads that express the *unc-119::gfp* transgene.

Sperm genes displayed elevated read counts in P-granule-depleted adult gonads compared to control adult gonads (Figure 3A and D). In hermaphrodite worms, spermatogenesis occurs during the L4 larval stage and ends at the L4-to-adult molt, at which time oogenesis begins (Hirsh *et al.* 1976). Consistent with this, transcripts from sperm-specific genes, such as the *msp* genes that encode MSPs or Major Sperm Proteins, accumulated in L4 gonads and then abruptly declined from control adult gonads. However, these transcripts remained elevated in P-granule-depleted adult gonads (Figure 3D and Figure S2C), supporting the suggestion from PCA analysis that P granules are necessary for germlines to progress from L4 to adulthood. These data are consistent with a recent study that also detected perdurance of sperm transcripts in P-granule RNAi Day 1 adult gonads (Campbell and Updike 2015).

Soma-specific genes showed several patterns of increased transcript accumulation in P-granule-depleted adult gonads (Figures 3E, S3, and S4). Some soma-specific transcripts accumulated in a pattern resembling sperm transcripts (transcripts detected in L4 gonads, down-regulated in control adult gonads, and present at elevated levels in P-granule-depleted adult gonads), including the alpha 2(IV) collagen mRNA, *let-2* (Figure 3E). Other transcripts, like *cebp-1*, which encodes a transcription factor important in axon regeneration in neurons (Yan *et al.* 2009), and *puf-9*, which encodes a Pumilio-related RNA-binding protein that is expressed in the hypodermis and neurons (Nolde *et al*. 2007), had very low read levels in control gonads but were significantly up-regulated in P-granule-depleted adult gonads at all stages (Figures 3E and S6A). In contrast to genes like *let-2, cebp-1*, and *puf-9*, some soma-specific transcripts were only significantly up-regulated in Day 2 Adult PG(-) UNC-119::GFP(+) gonads, including *aqp-2,* which encodes a membrane-bound aquaporin that is normally expressed in the excretory cell, muscle cells, motor neurons, and the hypodermis (Huang *et al.* 2007; Figure 3E). Although different somatic genes displayed different patterns of up-regulation (Figure 3E), as a whole, the levels of somatic gene transcripts progressively increased over time and with expression of the *unc-119::gfp* transgene (Figures 3A and S2D).

As an unbiased, genome-wide approach to determine different gene expression patterns among our samples, we also performed k-means clustering using the fold changes of RNAs in P-granule-depleted gonads versus their paired controls (see Materials and Methods). The 14,571 genes that showed a significant fold change in at least one of our comparisons grouped into eight clusters. The majority of ubiquitous genes, germline-enriched genes, and germline-specific genes grouped into Cluster 1, and the majority of sperm-specific genes grouped into Cluster 4 (Figures S3 and S4). This shows that these genes act similarly within their assigned gene category; that is, the majority of germline genes show similar responses to P-granule depletion, as do the majority of sperm genes. Somatic genes were spread out among the rest of the clusters (Clusters 2, 3, 5, 6, 7, and 8; Figure S3), supporting our observation that P-granule depletion causes several patterns of misregulation of soma-specific transcripts. Some somatic transcripts became more prevalent at progressively later stages (e.g., *cebp-1* in Cluster 2), while others were detected in L4 gonads, were reduced in control adult gonads, and persisted in PG(-) adult gonads (e.g., *let-2* in Cluster 6). We suspect that the somatic genes that become highly expressed in UNC-119::GFP(+) gonads (e.g., *aqp-2* and certain genes in Clusters 2, 3, and 6) are due to indirect, secondary effects of P-granule loss (see Discussion). Taken together, our transcript profiling shows that P granules are necessary to promote appropriate transcript accumulation patterns in gonads after the L4 larval stage.

### Single molecule RNA-FISH analysis confirms that somatic transcripts accumulate in P-granule-depleted gonads

Although our RNA-seq method is highly sensitive, it is unable to provide spatial information about transcript accumulation patterns within gonads. As an independent method to test if, when, and where soma-specific transcripts are up-regulated in P-granule-depleted germ cells, we performed single molecule RNA-FISH (smFISH) using probes to detect several soma-specific mRNAs (*cebp-1*, *aqp-2, let-2,* and *puf-9*) in control and P-granule-depleted gonads.

By RNA-seq, *cebp-1* transcripts were not detected in control L4 or adult gonads but were detected in P-granule-depleted gonads at all stages (Figure 3E). By smFISH, very few *cebp-1* transcripts were detected in control gonads at any stage (Figure 4A). We did detect some *cebp-1* transcripts around the nuclei of somatic gonad cells, including the very distal tip of the gonad, likely the somatic distal tip cell, in Day 1 and Day 2 Adult control gonads, but no signal was detected in the germline. Upon P-granule depletion, individual *cebp-1* mRNA molecules were present throughout the germ cell region of Day 1 and Day 2 Adult PG(-) UNC-119::GFP(-) gonads as well as Day 2 Adult PG(-) UNC-119::GFP(+) gonads (Figure 4A). Indeed, 58% of Day 1 Adult PG(-) UNC-119::GFP(-) gonads (n=24) and 100% of Day 2 Adult PG(-) UNC-119::GFP(+) gonads (n=13) had *cebp-1* transcripts throughout the germ cell region. Unlike *cebp-1* transcripts, which were detected in younger P-granule-depleted gonads by RNA-seq, *aqp-2* transcripts were only up-regulated in Day 2 Adult PG(-) UNC-119::GFP(+) gonads (Figure 3E). Similarly, by smFISH, *aqp-2* transcripts were only detected in Day 2 Adult PG(-) UNC-119::GFP(+) gonads (Figure 4B), and 100% of Day 2 Adult PG(-) UNC-119::GFP(+) gonads had *aqp-2* transcripts (n=23). Like *cebp-1* transcripts, *aqp-2* transcripts were distributed throughout the germline and sometimes concentrated in regions around nuclei (Figure 4B).

**Figure 4.**
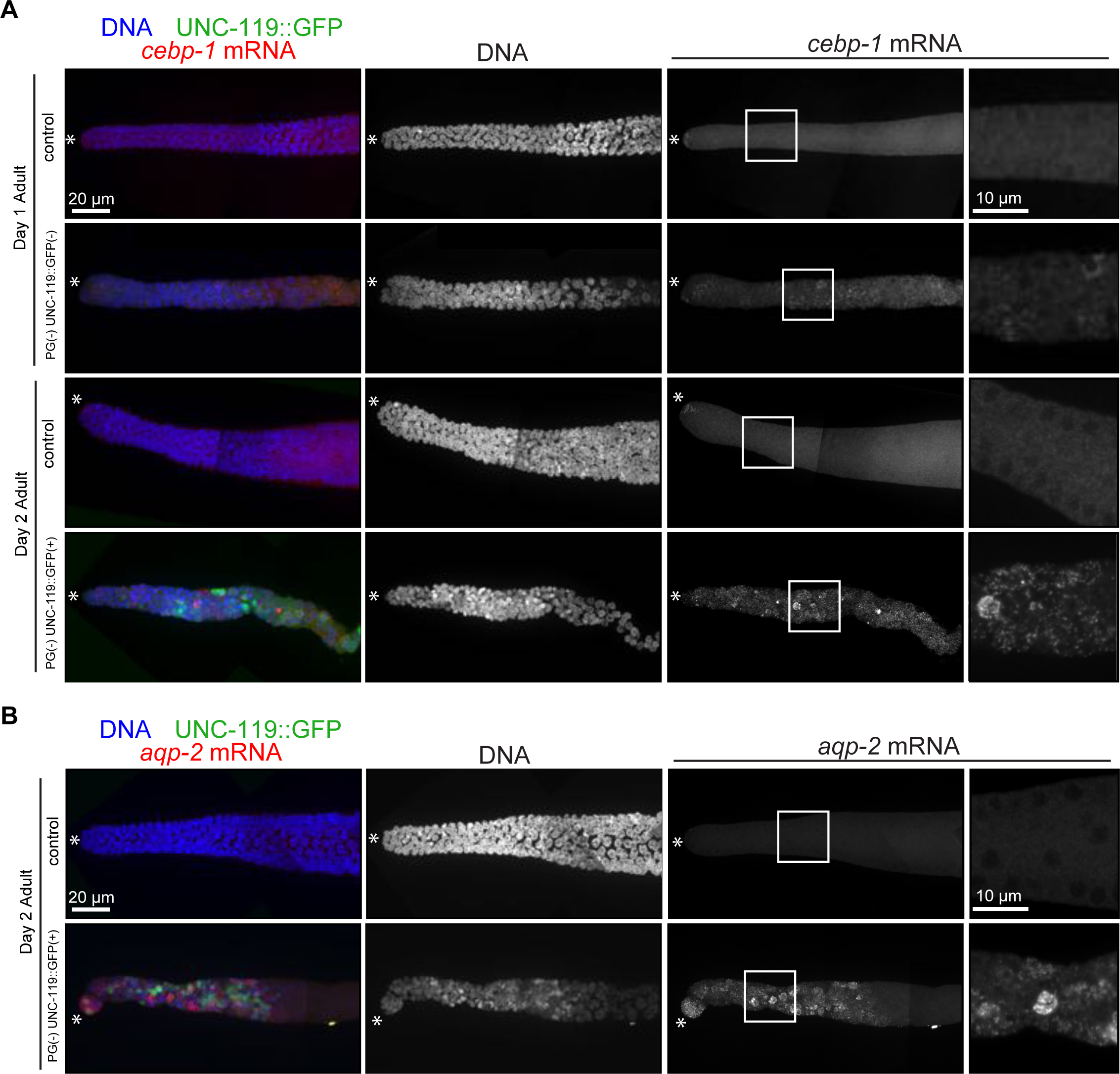
Soma-specific transcripts accumulate in P-granule-depleted germlines. **A)** Single molecule RNA-FISH (smFISH) analysis of *cebp-1* transcripts (red) in dissected Day 1 and Day 2 Adult control gonads, a Day 1 Adult PG(-) gonad, and a Day 2 Adult PG(-) UNC-119::GFP(+) gonad. **B)** smFISH analysis of *aqp-2* transcripts (red) in a Day 2 Adult control gonad and a Day 2 Adult PG(-) UNC-119::GFP(+) gonad. For each gonad in (A) and (B), UNC-119::GFP signal is in green and DAPI-stained DNA is in blue. The distal half of each gonad is shown as a z-stack projection, with the distal tip indicated by an asterisk. Boxed regions are expanded in the right-most panels. The zoomed-in regions are projections of 3 slices taken from the middle section of the gonad.

We also used smFISH to analyze *let-2* and *puf-9* transcripts. By RNA-seq, *let-2* transcripts in control gonads behaved in a similar manner to sperm transcripts; transcripts were detected in L4 gonads and decreased in adult gonads (Figure 3E). After P-granule depletion, *let-2* transcripts persisted in adults and were especially elevated in Day 2 Adult PG(-) UNC-119::GFP(+) gonads (Figure 3E). Interestingly, by smFISH, the *let-2* signal in both L4 control and L4 PG(-) gonads mostly originated from *let-2* transcripts concentrated in the very distal tip of the gonad, likely in the somatic distal tip cell and not in germ cells (Figure S5). In Day 2 Adult control gonads, *let-2* transcripts were mostly gone, with only a few transcripts remaining at the distal tip (Figure S5). In contrast, in 54% (n=36) of Day 2 Adult PG(-) UNC-119::GFP(+) gonads, *let-2* transcripts were present throughout the gonad, frequently concentrated in regions around the nuclei of germ cells (Figure S5). By RNA-seq, levels of *puf-9* transcripts were low in control gonads and elevated in both adult stages lacking P granules (Figure S6A). By smFISH, *puf-9* transcripts were also detected throughout the distal half of the germline in P-granule-depleted adult gonads: 71% of Day 1 Adult PG(-) (n=45) and 90% of Day 2 Adult PG(-) UNC-119::GFP(+) (n=19) (Figure S6B). Very few *puf-9* transcripts were detected in the distal half of control gonads. We did detect *puf-9* transcripts in the proximal half of both control and P-granule RNAi gonads. Taken together, our smFISH analysis corroborates our RNA-seq results and confirms that soma-specific transcripts are up-regulated in P-granule-depleted germ cells.

Our smFISH results shed light on the subset of soma-specific transcripts that are detected in L4 gonads, are reduced in control adult gonads, but persist in P-granule-depleted adult gonads (e.g., *let-2* transcripts in Figure 3E; Cluster 6 in Figures S3 and S4). Based on our RNA-seq data alone, two explanations are possible. One possibility is that these transcripts are expressed in germ cells at the L4 stage and then disappear in a P-granule-dependent manner in adult gonads, similar to sperm transcripts. An alternative possibility is that these transcripts reside in the somatic gonad and appear to be down-regulated in control adult gonads and up-regulated in P-granule-depleted adult gonads because of the altered ratio of somatic gonad cells to germ cells, i.e. a lower ratio in control adults and a higher ratio in P-granule-depleted adults with stunted germlines. Illustrating the latter possibility, our smFISH analysis detected *Iet-2* transcripts in the somatic gonad of L4s. However, *let-2* transcripts were later expressed in the germ cells of UNC-119::GFP(+) adults, along with many other somatic transcripts (e.g., *cebp-1*, *aqp-2*, and *puf-9*). Thus, even somatic gonad markers can become expressed in the germlines of reprogrammed adults. Further research is needed to identify the direct versus indirect effects that P granules have on individual genes.

### UNC-119::GFP(+) gonads turn on numerous genes involved in neuronal development

Most of the genes misregulated in Day 2 Adult PG(-) UNC-119::GFP(-) gonads were also misregulated in Day 2 Adult PG(-) UNC-119::GFP(+) gonads (Figure 2B). Day 2 Adult PG(-) UNC-119::GFP(+) gonads turned on many additional genes as well. Direct comparison of Day 2 Adult PG(-) UNC-119::GFP(+) and UNC-119::GFP(-) gonads identified 2,268 genes significantly up-regulated and 692 genes significantly down-regulated in UNC-119::GFP(+) gonads compared to UNC-119::GFP(-) gonads (Figure 5A). Gene Ontology analysis of the 2,268 up-regulated genes identified processes involved in neuron differentiation, synaptic transmission, and axon guidance, as well as a large number of genes involved in regulation of RNA metabolism (Figure 5B). Interestingly, we did not detect processes involved in development of other somatic cell types like hypodermis, intestine, or muscle, even though 60% of UNC-119::GFP(+) gonads also express the muscle myosin MYO-3 (Updike *et al*. 2014).

**Figure 5.**
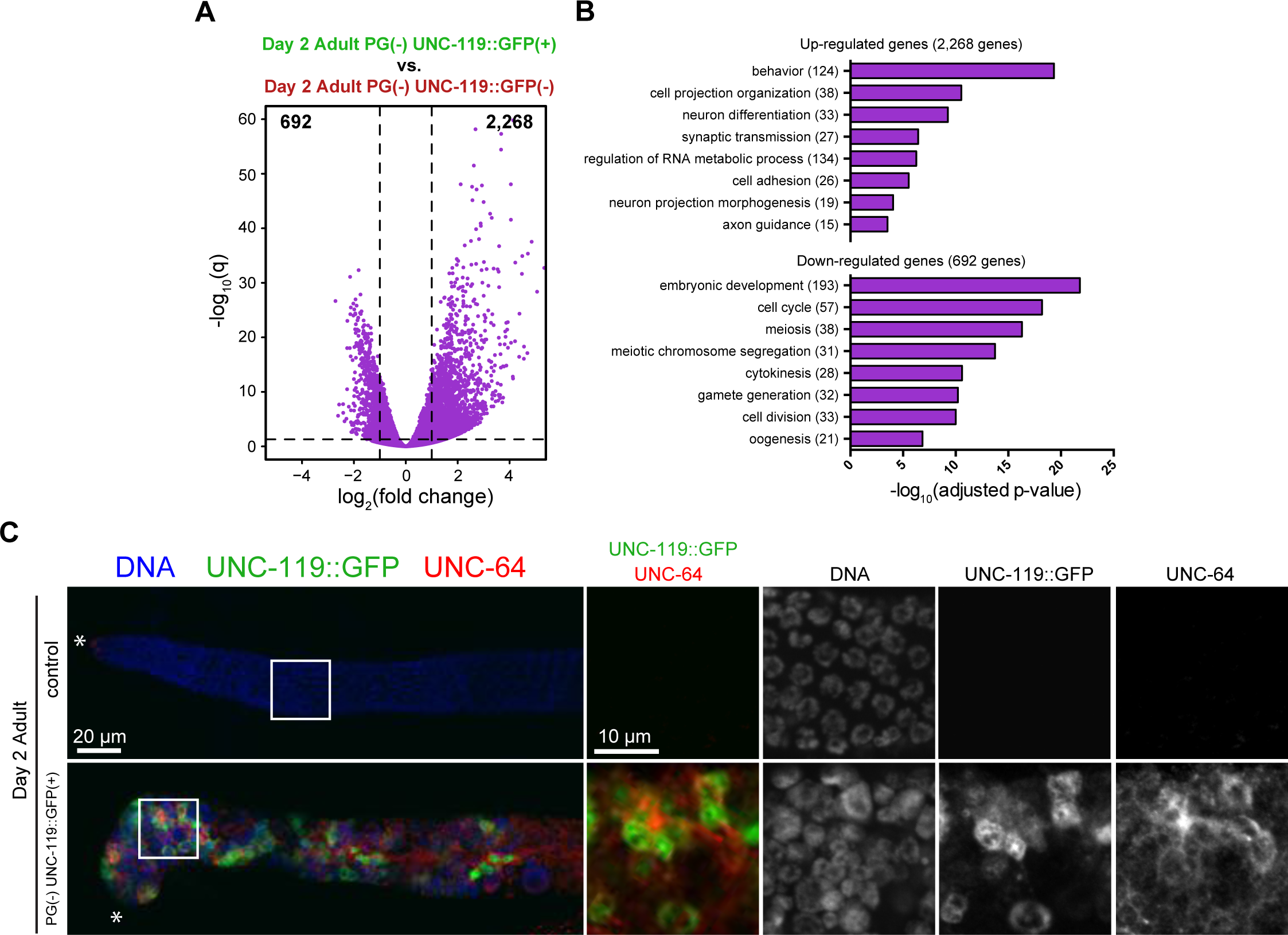
Day 2 Adult PG(-) UNC-119::GFP(+) gonads up-regulate genes involved in neuronal processes and down-regulate genes involved in germline processes. **A)** Volcano plot comparing Day 2 Adult PG(-) UNC-119::GFP(+) gonads to Day 2 Adult PG(-) UNC-119::GFP(-) gonads. Dashed lines indicate the significance cut-off of q=0.05 (horizontal line) and a 2-fold change in transcript level (vertical lines). The numbers of significantly misregulated genes in each quadrant are shown at the top of the plot. **B)** Gene Ontology analysis of the 2,268 up-regulated genes and 692 down-regulated genes in Day 2 Adult PG(-) UNC-119::GFP(+) gonads compared to Day 2 Adult PG(-) UNC-119::GFP(-) gonads. The Gene Ontology processes are organized based on their p-values, and the numbers of genes in each ontology category are shown in parentheses. **C)** Gonads from Day 2 Adult control and Day 2 Adult PG(-) UNC-119:GFP(+) worms were dissected and immunostained for UNC-64 (red), GFP (green), and DNA (blue). The distal half of each gonad is shown as a z-stack projection, with the distal tip indicated by an asterisk. Boxed regions are expanded in the right-most panels. The zoomed-in regions are projections of 3 slices taken from the middle section of the gonad.

As an independent test of whether neuronal genes other than the *unc-119::gfp* transgene are up-regulated in Day 2 Adult PG(-) UNC-119::GFP(+) gonads, we immunostained dissected gonads for the *C. elegans* syntaxin homolog UNC-64. UNC-64 is a component of the core synaptic vesicle fusion machinery and is expressed throughout the nervous system (Ogawa *et al.* 1998; Saifee *et al.* 1998). In our transcriptome data, *unc-64* transcripts were significantly up-regulated in Day 2 Adult PG(-) UNC-119::GFP(+) gonads compared to Day 2 Adult control and Day 2 Adult PG(-) UNC-119::GFP(-) gonads. In control gonads, some UNC-64 staining was detected in the somatic sheath and distal tip cell but not in the germ cell region (Figure 5C). In contrast, in Day 2 Adult PG(-) UNC-119::GFP(+) gonads, UNC-64 staining was observed throughout the germ cell region and stained in a filamentous pattern (Figure 5C), which may indicate cell membranes since syntaxin is a membrane-anchored receptor (Saifee *et al.* 1998). These data show that Day 2 Adult PG(-) UNC-119::GFP(+) gonads up-regulate many genes involved in neuronal development, in addition to the *unc-119::gfp* transgene.

### UNC-119::GFP(+) germ cells lose germline identity

Gene Ontology analysis of the 692 genes down-regulated in Day 2 Adult PG(-) UNC-119::GFP(+) versus UNC-119::GFP(-) gonads revealed processes involved in the cell cycle, meiosis, and chromosome segregation (Figure 5B). Interestingly, a large number of down-regulated genes were categorized as important for embryonic development, including genes whose gene products are maternally loaded into developing oocytes to support and guide early embryogenesis (e.g., *mex-5*, *mex-6*, *pie-1*, *nos-1*). To test if UNC-119::GFP(+) cells lose germ cell identity upon acquisition of a neuronal fate, we stained Day 2 Adult PG(-) UNC-119::GFP(+) gonads for the germline proteins REC-8 and HTP-3 and performed smFISH to detect *lag-1* transcripts. REC-8 is a cohesin found in the mitotic stem cell region of germlines (Hansen *et al*. 2004), and HTP-3 is an integral component of the synaptonemal complex present in the meiotic region (Severson *et al*. 2009; Figure 6A and 6C). LAG-1 is a transcription factor that works downstream of Notch signaling to control germ cell entry into meiosis (Crittenden *et al*. 2003). In Day 2 Adult PG(-) UNC-119::GFP(+) gonads, we observed a mutually exclusive pattern of UNC-119::GFP and REC-8 staining; cells that expressed UNC-119::GFP lacked REC-8 and vice versa (Figure 6A and B). This was also the case for HTP-3 (Figure 6C and D) and *lag-1* transcripts (Figure 6E). This demonstrates the antagonistic relationship between somatic and germline fates and highlights the importance of P granules in maintaining germ cell identity.

**Figure 6.**
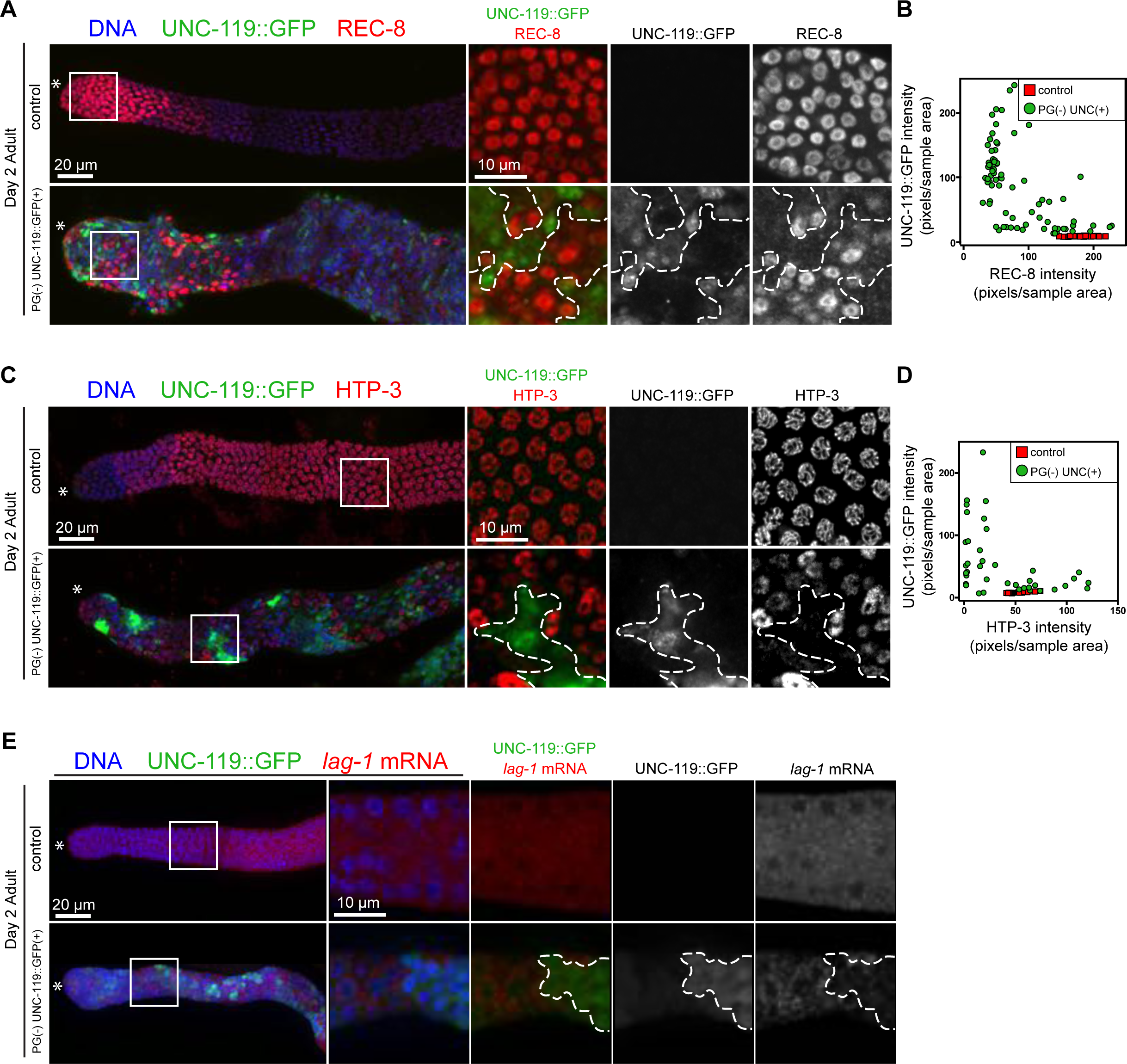
Acquisition of neuronal fate in P-granule-depleted germ cells is accompanied by loss of germ cell markers. **A)** Gonads from Day 2 Adult control and Day 2 Adult PG(-) UNC-119::GFP(+) worms were dissected and immunostained for REC-8 (red), GFP (green), and DNA (blue). **B)** Quantification of UNC-119::GFP signal relative to REC-8 signal in germ cells from control and P-granule-depleted gonads. **C)** Same as in (A) except gonads were stained for HTP-3 (red). **D)** Same as in (B) except gonads were quantified for UNC-119::GFP signal relative to HTP-3 signal. **E)** Single molecule RNA-FISH (smFISH) analysis of *lag-1* transcripts (red) in a dissected Day 2 Adult control gonad and a Day 2 Adult P granule(-), UNC-119::GFP(+) gonad. For (A), (C), and (E), the distal half of each gonad is shown as a z-stack projection, with the distal tip indicated by an asterisk. Boxed regions are expanded in the right-most panels. The zoomed-in regions are projections of 3 slices taken from the middle section of the gonad, except for the control HTP-3 gonad (C), which shows the top section of the gonad. In the expanded panels in (C), the red channel of the PG(-) gonad was increased to better show relative HTP-3 levels.

Taken together, our transcript profiling, immunostaining, and smFISH experiments demonstrate that changes to the transcriptome and proteome are occurring in at least a fraction of the germ cells in P-granule-depleted gonads. These changes reflect both the loss of germline identity (e.g., loss of REC-8, HTP-3, and *lag-1* mRNA) and acquisition of somatic identity (e.g., expression of the *unc-119::gfp* transgene and UNC-64).

### The germlines from *pgl-1* and *glh-1* single mutants also express somatic markers

Our P-granule RNAi construct simultaneously targets four important P-granule genes: *pgl-1*, *pgl-3*, *glh-1*, and *glh-4*. We tested if loss of one or two P-granule components is sufficient to cause germline expression of somatic genes by crossing the *unc-119::gfp* transgene into *glh-1* single and *glh-1 glh-4* double mutant backgrounds and by performing single *pgl-1* RNAi or double *pgl-1; pgl-3* RNAi on *unc-119::gfp* transgenic worms. We detected UNC-119::GFP expression in all backgrounds tested (Figures 7A and S7). Additionally, of the germlines that expressed UNC-119::GFP, a fraction also expressed the muscle myosin MYO-3 (Figures 7A and S7). To determine if *pgl-1* and *glh-1* mutant gonads misregulate a similar set of genes, we performed transcript profiling of gonads collected from sterile Day 1 Adult *pgl-1* mutants and sterile Day 1 Adult *glh-1* mutants; neither mutant contained the *unc-119::gfp* transgene. Comparison to wild-type gonads revealed more transcript accumulation changes in *pgl-1* mutants than in *glh-1* mutants; *pgl-1* single mutants up-regulated 2,894 genes and down-regulated 273 genes, while *glh-1* single mutants up-regulated 626 genes and down-regulated 110 genes (Figure 7B). The majority of the genes up-regulated in *glh-1* gonads were also up-regulated in *pgl-1* gonads (592 out of 626 genes, 94%), suggesting that PGL-1 and GLH-1 may work together to control certain transcripts (Figure 7C). The majority of genes up-regulated in both *pgl-1* and *glh-1* mutant gonads were also up-regulated in Adult P-granule RNAi gonads (Figures S3 and S4; data not shown). These results suggest that PGL-1 and GLH-1 are both critical P-granule factors and that both are necessary to prevent somatic development within the germline.

**Figure 7.**
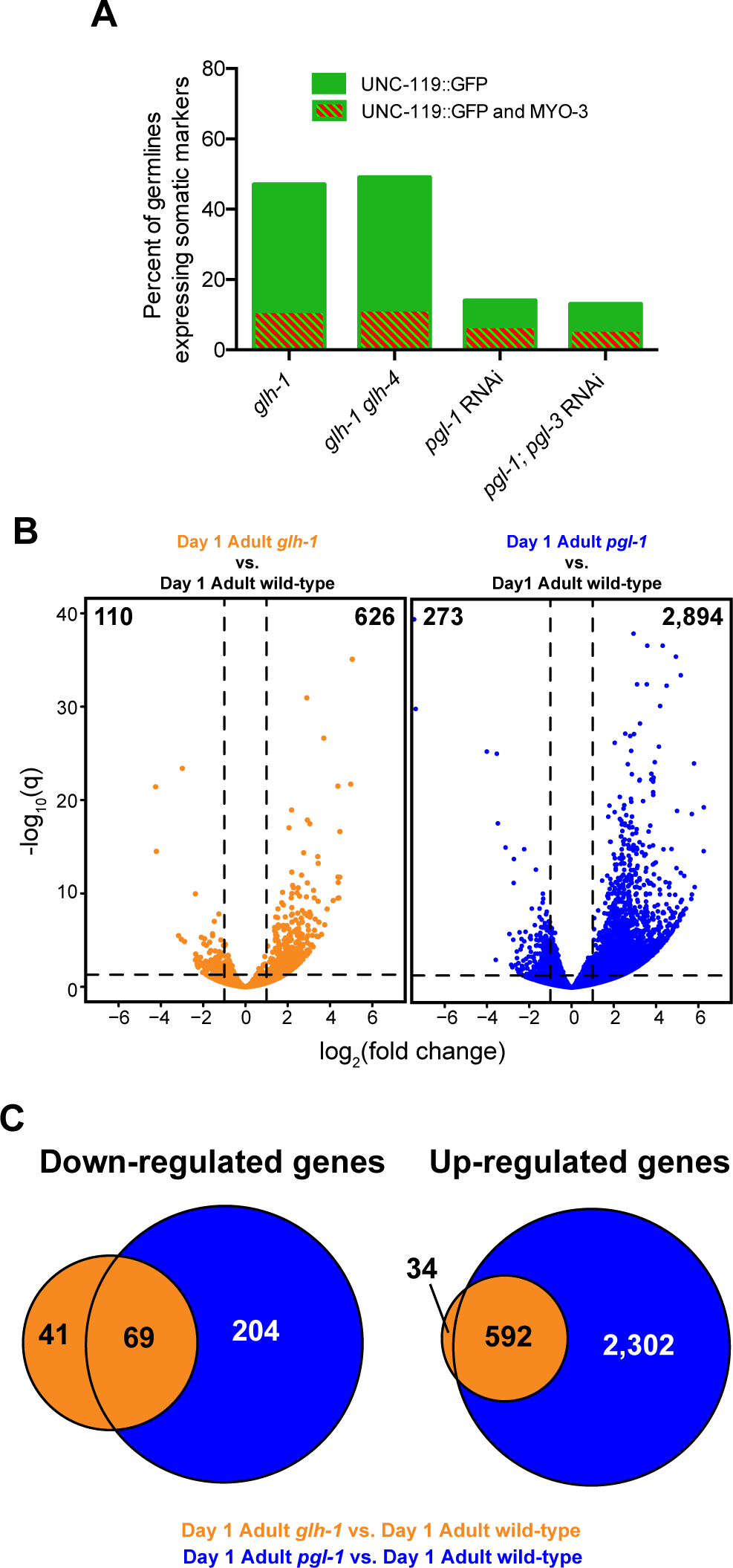
Germlines lacking PGL-1 or GLH-1 also misexpress somatic markers. **A)** Percent of germlines expressing the neuronal marker UNC-119::GFP (green bars) in *glh-1* single mutants, *glh-1 glh-4* double mutants, *pgl-1* single RNAi, and *pgl-1; pgl-3* double RNAi animals containing the *unc-119::gfp* transgene. Of the germlines that expressed UNC-119::GFP, a fraction also expressed the muscle marker MYO-3 (green and red striped bars). **B)** Volcano plots comparing *glh-1* (gold) or *pgl-1* (blue) single mutant gonads to wild-type gonads. All animals lacked the *unc-119::gfp* transgene. Dashed lines indicate the significance cut-off of q=0.05 (horizontal line) and a 2-fold change in transcript levels (vertical lines). The numbers of significantly misregulated genes in each quadrant are shown at the top of the plot. **C)** Venn diagrams showing the overlap of genes down‐ or up-regulated in *glh-1* and *pgl-1* single mutant gonads.

## DISCUSSION

Although germ granules were first observed over 100 years ago, their functions are only starting to be understood at the molecular level. *C. elegans* germ granules or “P granules” are necessary for germline development and fertility (Kawasaki *et al*. 1998; Kawasaki *et al*. 2004; Spike *et al*. 2008) and protect the germline from expressing somatic fate markers (Updike *et al.* 2014). In this study, we analyzed the progressive transcript changes that occur in P-granule-depleted germlines. We treated worms with P-granule RNAi for two generations and performed transcriptome analysis of dissected gonads from the second-generation sterile hermaphrodites. In worms that lack P granules, germlines appear relatively healthy at the L4 stage but deteriorate in adults. Germline deterioration in adults is accompanied by and perhaps caused by aberrant persistence of sperm-specific transcripts after the sperm-production period in L4, as previously observed (Campbell and Updike 2015), and elevated levels of soma-specific transcripts. The persistence of sperm-specific transcripts into adulthood suggests that P granules promote progression of germ cells from the L4 larval stage to adulthood, and the healthy appearance and normal RNA profile of L4 germlines suggest that P granules are not necessary for germline development from embryogenesis to L4. In adults, the majority of P-granule-depleted germlines become reduced in size and disorganized, while a minority of germlines appear to proliferate; both categories of germlines can contain cells that have lost their previous germ cell fate and appear to be on a neuronal developmental path. Taken together, these findings reveal that *C. elegans* germ granules are major regulators of the germline transcriptome and prevent germ cells from undergoing somatic development, possibly by preventing the expression of somatic factors in the germline (Figure 8). The mechanism by which P granules perform these functions remains to be determined.

**Figure 8.**
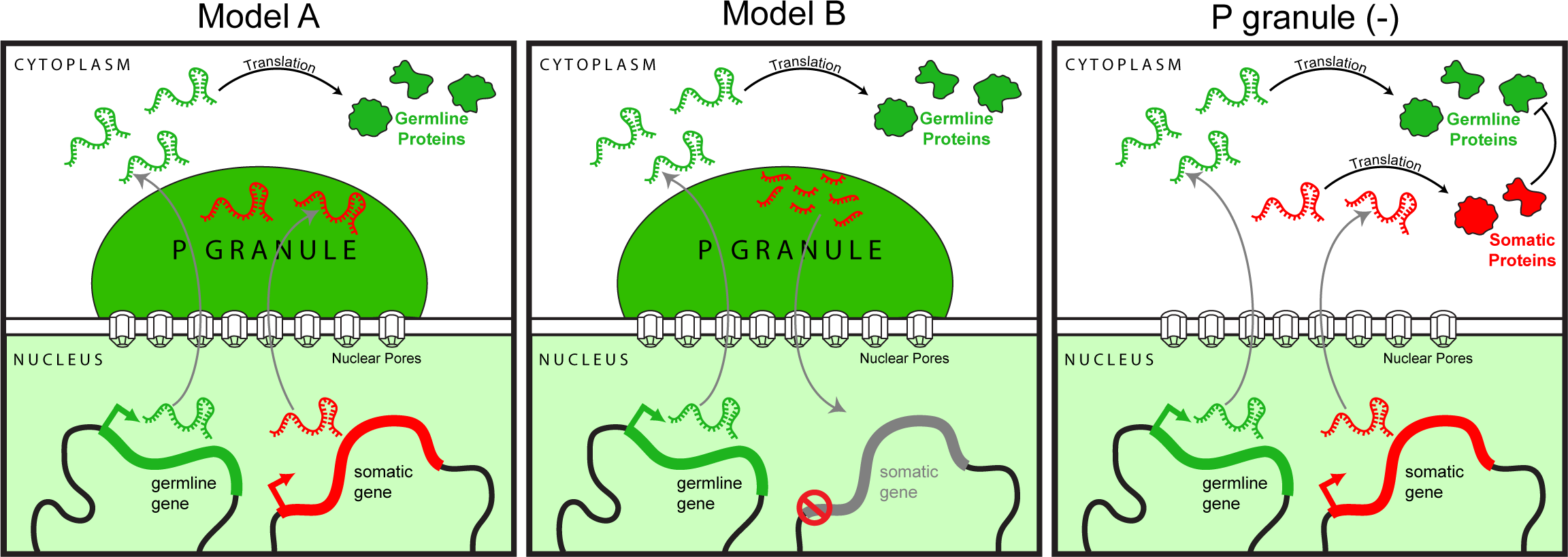
Models of P-granule function. P granules reside over nuclear pores, where they intercept stochastically misexpressed soma-specific mRNAs (red mRNAs), while allowing germline-appropriate mRNAs (green mRNAs) to transit to the cytoplasm for translation. In Model A, P granules prevent the translation of somatic mRNAs. In Model B, P granules slice somatic mRNAs into small RNAs, which in turn induce heritable silencing of somatic genes. In the absence of P granules (right panel), somatic mRNAs accumulate and are translated into proteins (red) that specify somatic identity and antagonize germline identity.

P granules reside on the outer periphery of germ cell nuclei and overlie nuclear pores (Pitt *et al* 2000; Sheth *et al*. 2010; Updike *et al*. 2011). It has been proposed that as mRNAs exit the nucleus, P granules intercept and perhaps process these transcripts, thus influencing the transcriptome and proteome of the germline (Kasper *et al*. 2014; Figure 8). Indeed, some germline transcripts have been shown to at least transiently reside in P granules (Schisa *et al*. 2001; Sheth *et al*. 2010). Recently, PGL-1 was shown to dimerize and to have guanosine-specific RNase activity (Aoki *et al*. 2016). This raises the possibility that PGL-1 directly influences the transcriptome by cleaving its associated mRNAs, although PGL-1’s catalytic activity is not essential for its role in fertility (Aoki *et al*. 2016). If PGL-1 indeed has RNase activity *in vivo*, then an attractive scenario is that it recognizes mRNAs inappropriate for germline development and cleaves them, while allowing germline-appropriate mRNAs to pass through P granules and be translated in the cytoplasm (Figure 8). Alternatively, PGL-1 and the paralogous protein PGL-3 may not have a preference for transcript type. Indeed, PGL-3 was recently shown to bind mRNA in extracts and *in vitro* in a sequence-independent manner (Saha *et al*. 2016). One possibility is that P granules intercept all types of transcripts, including germline-appropriate transcripts, but effectively prevent low levels of stochastically expressed soma-specific transcripts from producing levels of proteins that would threaten germ cell identity.

Our study extends a recent analysis of transcripts from Day 1 adult P-granule RNAi gonads (Campbell and Updike 2015) and documents the progressive appearance of somatic transcripts that underlies the germ-toward-soma reprogramming we previously observed in older P-granule RNAi adults (Updike *et al*. 2014). Campbell and Updike documented perdurance of sperm-specific transcripts in Day 1 adult P-granule RNAi gonads, as we describe in both Day 1 and Day 2 P-granule RNAi adults, but they did not detect increased accumulation of soma-specific transcripts in their samples. The Day 1 adult worms sampled by Campbell and Updike experienced weaker RNAi (based on RNA read counts from RNAi target genes) and contained healthier germlines than in our experiments. Thus, the defects observed by Campbell and Updike are probably the earliest defects and likely include the primary defects caused by P-granule depletion. Our stronger RNAi and additional analysis of Day 2 adults with progressively reprogrammed germlines identified additional consequences, probably many secondary consequences, of P-granule depletion. Based on their observations, Campbell and Updike concluded that P granules likely repress translation of soma-specific mRNAs that are already present at low levels in the germline (Figure 8, Model A). Our data suggest that P granules additionally affect the levels of soma-specific mRNAs in the germline, possibly by degrading soma-specific mRNAs and/or repressing transcription of soma-specific genes (Kasper *et al*. 2014; Figure 8, Model B).

Consistent with P granules functioning as regulators of mRNA accumulation in the germline, many RNAi factors localize to P granules, including Dicer/DCR-1, the RNA-dependent RNA polymerase EGO-1, and the argonaute proteins PRG-1 and CSR-1 (Batista *et al*. 2008; Claycomb *et al*. 2009). PRG-1 is a Piwi-related argonaute that prevents the expression of foreign genetic elements, including transposon and transgene mRNAs. PRG-1 recognizes these mRNAs through imperfect base-pairing with its associated 21U-RNAs, resulting in heritable silencing of transposon and transgene loci (Batista *et al*. 2008; Ashe *et al*. 2012; Bagijn *et al*. 2012; Shirayama *et al*. 2012; Lee *et al*. 2012). To maintain germline identity, PRG-1 may work with P-granule proteins to recognize and degrade soma-specific transcripts identified as “foreign”. This information could then be relayed back into the nucleus to repress inappropriately activated soma-specific genes through epigenetic mechanisms (Kasper *et al*. 2014; Figure 8). CSR-1, another P-granule associated argonaute, recognizes its targets through associated 22G-RNAs that base-pair with germline-expressed mRNAs including those important for embryogenesis (Claycomb *et al*. 2009). Interestingly, CSR-1 was recently found to tune the levels of certain germline mRNAs in a manner that is dependent on its slicer activity (Gerson-Gurwitz *et al*. 2016). Importantly, depletion of CSR-1 and depletion of P granules cause similar transcript accumulation changes (Campbell and Updike 2015), strengthening the link between argonautes and P granules in regulating the germline transcriptome.

The development of different neuronal subtypes can be induced in the *C. elegans* germline by simultaneous removal of certain chromatin factors and forced expression of master neuronal transcription factors (Tursun *et al*. 2011; Patel *et al*. 2012), highlighting the plasticity of the germline. Notably, other genetic perturbations have been shown to cause *C. elegans* germ cells to express neuronal and muscle markers, without the need for driving transcription factors. These include loss of the H3K4 methyltransferase SET-2, loss of the nuclear argonaute HRDE-1, and concomitant loss of the NuRD complex component LET-418 and the H3K4 demethylase SPR-5 (Robert *et al*. 2014; Kaser-Pebernard *et al*. 2014). Because these factors are chromatin-based, they are likely to have direct effects on gene transcription. One possibility is that these factors work downstream of P granules and interpret the RNA-processing events that occur within P granules. Comparing the transcriptome of gonads lacking P granules to the transcriptomes of *set-2* or *let-418; spr-5* mutant gonads may provide insight into how these regulatory mechanisms operate and influence each other to protect germ cell identity.

Our Gene Ontology analysis of genes uniquely up-regulated in P-granule-depleted gonads from Day 2 adults expressing the *unc-119::gfp* neuronal transgene revealed processes in neuronal development. Interestingly, processes involved in muscle development and function were not observed, despite the fact that 60% of UNC-119::GFP(+) gonads also express the muscle myosin MYO-3 (Updike *et al*. 2014). Perhaps many fewer P-granule-depleted germ cells up-regulate genes involved in muscle development than up-regulate genes involved in neuronal development, which would bias our transcriptome analysis in favor of genes involved in neuron-related processes.

The expression of many neuronal genes in P-granule-depleted Day 2 adult germlines is likely to be a secondary effect of P-granule loss. The burning challenges are to identify the primary effect(s) of P-granule loss, and to determine why P-granule-depleted germ cells appear to favor a neuronal path. One possibility is that these germ cells are reverting to a “default” neuronal differentiation program, which has been observed in the contexts of dissociated *Xenopus* embryos and mammalian embryonic stem cells (ESCs) grown in culture (Sato and Sargent 1989; Tropepe *et al*. 2001). To acquire neuronal fate through this “default” program, ESCs must be cultured at low-density and in growth factor-free conditions, suggesting an inherent propensity of ESCs to differentiate into neurons (Smukler *et al*. 2006). Recently, the transcription factor BLIMP1 was found to be essential to reprogram human induced pluripotent stem cells (hiPSCs) into primordial germ cell (PGC)-like cells (Sasaki *et al*. 2015). Interestingly, when Sasaki *et al*. attempted to derive PGC-like cells from *BLIMP1-/-* hiPSCs, essential PGC genes were not expressed and genes involved in neuronal differentiation were up-regulated instead. These and our findings suggest that neuronal fate is the preferred somatic fate of cells that lose their pluripotent stem cell or germline identity. Future work should investigate if *C. elegans* germlines that lack P granules revert to a “default” neuronal program and if so, how the neuronal program becomes activated and why it becomes activated in adults and not larval germlines.

The breadth of ribonucleoprotein complexes and the regulatory circuits they form within the *C. elegans* germline highlight the complex nature of this tissue. In order to fully understand P-granule function, a thorough analysis of individual factors like PGL-1 and GLH-1 will need to be performed, especially at the biochemical level. Additionally, identification of P-granule-associated mRNAs may provide insight into how P granules prevent somatic development in the germline. Finally, because many P-granule proteins are conserved in other species, it is important to test if germ granules in other organisms including vertebrates prevent somatic development of germ cells.

## AUTHOR CONTRIBUTIONS

A.K.K., T.E., and S.S. designed the research. A.K.K. performed the RNA-seq and immunostaining experiments. T.E. performed the mutant analyses and smFISH experiments. A.K.K. and A.R. performed the RNA-seq bioinformatic analysis. A.K.K., T.E. and S.S. wrote the paper.

## ACKNOWLEDGEMENTS

We thank Geraldine Seydoux, David Greenstein, and Judith Kimble for advice on smFISH, Judith Kimble for the *lag-1* smFISH probe, Mike Nonet for the UNC-64 antibody, Dustin Updike for the *pgl-1; pgl-3* double RNAi construct, Ben Abrams at the UCSC Life Sciences Microscopy Facility for training, and past and current members of the Strome lab for helpful discussions. This work used the Vincent J. Coates Genomics Sequencing Laboratory at UC Berkeley, supported by NIH S10 Instrumentation Grants (S10RR029668, S10RR027303, and S10OD018174). Some strains used in this study were provided by the CGC, which is funded by the NIH Office of Research Infrastructure Programs (P40 OD010440). This work was supported by a CIRM predoctoral fellowship (TG2-01157) to A.K.K. and an NIH R01 grant (GM34059) and an Ellison Senior Scholar in Aging Award (AG-SS-3098-12) to S.S.

## REFERENCES

Alder, M. N., S. Dames, J. Gaudet, and S. E. Mango, 2003 Gene silencing in *Caenorhabditis elegans* by transitive RNA interference. RNA 9: 25–32.

Aoki S. T., A. M. Kershner, C. A. Bingman, M. Wickens, and J. Kimble, 2016 PGL germ granule assembly protein is a base-specific, single-stranded RNase. Proc. Natl. Acad. Sci. USA 113: 1279–1284.

Ashe A., A. Sapetschnig, E. M. Weick, J. Mitchell, M. P. Bagijn et al., 2012 piRNAs can trigger a multigenerational epigenetic memory in the germline of *C. elegans*. Cell 150: 88–99.

Bagijn M. P., L. D. Goldstein, A. Sapetschnig, E. Weick, S. Bouasker et al., 2013 Function, targets, and evolution of *Caenorhabditis elegans* piRNAs. Science 337: 574–578.

Batista P. J., J. G. Ruby, J. M. Claycomb, R. Chiang, N. Fahlgren et al., 2008 PRG-1 and 21U‐ RNAs interact to form the piRNA complex required for fertility in *C. elegans*. Mol. Cell 31: 67–78.

Baugh L. R., A. A. Hill, D. K. Slonim, E. L. Brown, and Hunter C. P., 2003 Composition and dynamics of the *Caenorhabditis elegans* early embryonic transcriptome. Development 130: 889–900.

Brenner S., 1974 The genetics of *Caenorhabditis elegans*. Genetics 77: 71–94.

Campbell A. C. and D. L. Updike, 2015 CSR-1 and P granules suppress sperm-specific transcription in the *C. elegans* germline. Development 142: 1745–1755.

Chuma S., M. Hosokawa, T. Tanaka, and N. Nakatsuji, 2009 Ultrastructural characterization of spermatogenesis and its evolutionary conservation in the germline: Germinal granules in mammals. Mol. Cell. Endocrinol. 306: 17–23.

Claycomb J. M., P. J. Batista, K. M. Pang, W. Gu, J. J. Vasale et al., 2009 The argonaute CSR-1 and its 22G-RNA cofactors are required for holocentric chromosome segregation. Cell 139: 123–134.

Crittenden S. L., C. R. Eckmann, L. Wang, D. S. Bernstein, M. Wickens et al., 2003 Regulation of the mitosis/meiosis decision in the *Caenorhabditis elegans* germline. Philos. Trans. R. Soc. L. B. Biol. Sci. 358: 1359–1362.

Edelstein A. D., M. A. Tsuchida, N. Amodaj, H. Pinkard, R. D. Vale et al., 2014 Advanced methods of microscope control using μManager software. J. Biol. Methods 1: 10.

Gallo C. M., J. T. Wang, F. Motegi, and G. Seydoux, 2010 Cytoplasmic partitioning of P granule components is not required to specify the germline in *C. elegans*. Science 330: 1685–1689.

Gerson-Gurwitz A., S. Wang, S. Sathe, R. Green, G. W. Yeo et al., 2016 A Small RNA-Catalytic Argonaute Pathway Tunes Germline Transcript Levels to Ensure Embryonic Divisions. Cell 165: 396–409.

Gruidl M. E., P. A. Smith, K. A. Kuznicki, J. S. McCrone, J. Kirchner et al., 1996 Multiple potential germ-line helicases are components of the germ-line-specific P granules of *Caenorhabditis elegans*. Proc. Natl. Acad. Sci. USA 93: 13837–13842.

Hansen D., E. J. A. Hubbard, and T. Schedl, 2004 Multi-pathway control of the proliferation versus meiotic development decision in the *Caenorhabditis elegans* germline. Dev. Biol. 268: 342–357.

Hirsh D., D. Oppenheim, and M. Klass, 1976 Development of the reproductive system of *Caenorhabditis elegans*. Dev. Biol. 49: 200–219.

Huang C. G., T. Lamitina, P. Agre, and K. Strange, 2007 Functional analysis of the aquaporin gene family in *Caenorhabditis elegans*. Am. J. Physiol. Cell Physiol. 292: C1867–C1873.

Kaser-Pebernard S., F. Muller, and C. Wicky, 2014 LET-418/Mi2 and SPR-5/LSD1 cooperatively prevent somatic reprogramming of *C. elegans* germline stem cells. Stem Cell Reports 2: 547–559.

Kasper D. M., K. E. Gardner, and V. Reinke, 2014 Homeland security in the *C. elegans* germ line: Insights into the biogenesis and function of piRNAs. Epigenetics 9: 62–74.

Kawasaki I., Y. H. Shim, J. Kirchner, J. Kaminker, W. B. Wood et al., 1998 PGL-1, a predicted RNA-binding component of germ granules, is essential for fertility in *C. elegans*. Cell 94: 635–645.

Kawasaki I., A. Amiri, Y. Fan, N. Meyer, S. Dunkelbarger et al., 2004 The PGL family proteins associate with germ granules and function redundantly in *Caenorhabditis elegans* germline development. Genetics 167: 645–661.

Kim D., G. Pertea, C. Trapnell, H. Pimentel, R. Kelley et al., 2013 TopHat2: accurate alignment of transcriptomes in the presence of insertions, deletions and gene fusions. Genome Biol. 14: R36.

Knaut H., F. Pelegri, K. Bohmann, H. Schwarz, and C. Nüsslein-Volhard, 2000 Zebrafish vasa RNA but not its protein is a component of the germ plasm and segregates asymmetrically before germline specification. J. Cell Biol. 149: 875–888.

Knutson A. K., A. Rechtsteiner, and S. Strome, 2016 Reevaluation of whether a soma-to-germ-line transformation extends lifespan in *Caenorhabditis elegans*. Proc. Natl. Acad. Sci. USA 113: 3591–3596.

Kuznicki K. A., P. A. Smith, W. M. Leung-Chiu, A. O. Estevez, H. C. Scott et al., 2000 Combinatorial RNA interference indicates GLH-4 can compensate for GLH-1; these two P granule components are critical for fertility in *C. elegans*. Development 127: 2907–2916.

Lee H., W. Gu, M. Shirayama, E. Youngman, D. Conte et al., 2012 *C. elegans* piRNAs mediate the genome-wide surveillance of germline transcripts. Cell 150: 78–87.

Lee C., E. B. Sorensen, T. R. Lynch, and J. Kimble, 2016 *C. elegans* GLP-1/Notch activates transcription in a probability gradient across the germline stem cell pool. eLife 5: e18370.

Lesch B. J. and D. C. Page, 2012 Genetics of germ cell development. Nat. Rev. Genet. 13: 781–794.

Liu T., A. Rechtsteiner, T. A. Egelhofer, A. Vielle, I. Latorre et al., 2011 Broad chromosomal domains of histone modification patterns in *C. elegans*. Genome Res. 21: 227–236.

Love M. I., W. Huber, and S. Anders, 2014 Moderated estimation of fold change and dispersion for RNA-seq data with DESeq2. Genome Biol. 15: 550.

MacQueen A. J., C. M. Phillips, N. Bhalla, P. Weiser, A. M. Villeneuve et al., 2005 Chromosome sites play dual roles to establish homologous synapsis during meiosis in *C. elegans*. Cell 123: 1037–1050.

Meissner B., A. Warner, K. Wong, N. Dube, A. Lorch et al., 2009 An integrated strategy to study muscle development and myofilament structure in *Caenorhabditis elegans*. PLoS Genet. 5: e1000537

Nolde M., N. Saka, K. L. Reinert, and F. J. Slack, 2007 The *C. elegans pumilio* homolog, *puf-9*, is required for the 3’UTR mediated repression of the *let-7* microRNA target gene, *hbl-1*. Dev. Biol. 305: 551–563.

Ogawa H., S. I. Harada, T. Sassa, H. Yamamoto, and R. Hosono, 1998 Functional properties of the unc-64 gene encoding a *Caenorhabditis elegans* syntaxin. J. Biol. Chem. 273: 2192–2198.

Patel T., B. Tursun, D. P. Rahe, and O. Hobert, 2012 Removal of Polycomb Repressive Complex 2 makes *C. elegans* germ cells susceptible to direct conversion into specific somatic cell types. Cell Rep. 2: 1178–1186.

Pitt J. N., J. A. Schisa, and J. R. Priess, 2000 P granules in the germ cells of *Caenorhabditis elegans* adults are associated with clusters of nuclear pores and contain RNA. Dev. Biol. 219: 315–333.

Priess J. R., and J. N. Thomson, 1987 Cellular interactions in early *C. elegans* embryos. Cell 48: 241–250.

Rechtsteiner A., S. Ercan, T. Takasaki, T. M. Phippen, T. A. Egelhofer et al., 2010 The histone H3K36 methyltransferase MES-4 acts epigenetically to transmit the memory of germline gene expression to progeny. PLoS Genet. 6: e1001091.

Reinke V., I. S. Gil, S. Ward, and K. Kazmer, 2004 Genome-wide germline-enriched and sex‐ biased expression profiles in *Caenorhabditis elegans*. Development 131: 311–323.

Robert V. J., M. G. Mercier, C. Bedet, S Janczarski, J. Merlet et al., 2014 The SET-2/SET1 histone H3K4 methyltransferase maintains pluripotency in the *Caenorhabditis elegans* germline. Cell Rep. 9: 443–450.

Saha S., C. A. Weber, M. Nousch, O. Adame-Arana, C. Hoege et al., 2016 Polar positioning of phase-separated liquid compartments in cells regulated by an mRNA competition mechanism. Cell 166: 1572–1584.

Saifee O., L. Wei, and M. L. Nonet, 1998 The *Caenorhabditis elegans* unc-64 locus encodes a syntaxin that interacts genetically with synaptobrevin. Mol Biol Cell 9: 1235–1252.

Sasaki K., S. Yokobayashi, T. Nakamura, I. Okamoto, Y. Yabuta et al., 2015 Robust in vitro induction of human germ cell fate from pluripotent stem cells. Cell Stem Cell 17: 178–194.

Sato S. M. and T. D. Sargent, 1989 Development of neural inducing capacity in dissociated *Xenopus* embryos. Dev. Biol. 134: 263–266.

Schisa J. A., J. N. Pitt, and J. R. Priess, 2001 Analysis of RNA associated with P granules in germ cells of *C. elegans* adults. Development 128: 1287–1298.

Severson A. F., L. Ling, V. van Zuylen, and B. J. Meyer, 2009 The axial element protein HTP-3 promotes cohesin loading and meiotic axis assembly in *C. elegans* to implement the meiotic program of chromosome segregation. Genes Dev. 23: 1763–1778.

Sheth U., J. Pitt, S. Dennis, and J. R. Priess, 2010 Perinuclear P granules are the principal sites of mRNA export in adult *C. elegans* germ cells. Development 137: 1305–1314.

Shirayama M., M. Seth, H. C. Lee, W. Gu, T. Ishidate et al., 2012 piRNAs initiate an epigenetic memory of nonself RNA in the *C. elegans* germline. Cell 150: 65–77.

Smukler S. R., S. B. Runciman, S. Xu, and D. van der Kooy, 2006 Embryonic stem cells assume a primitive neural stem cell fate in the absence of extrinsic influences. J. Cell Biol. 172: 79–90.

Spike C., N. Meyer, E. Racen, A. Orsborn, J. Kirchner et al., 2008 Genetic analysis of the *Caenorhabditis elegans* GLH family of P-granule proteins. Genetics 178: 1973–1987.

Strome S. and W. B. Wood, 1983 Generation of asymmetry and segregation of germ-line granules in early *C. elegans* embryos. Cell 35: 15–25.

Strome S. and D. Updike, 2015 Specifying and protecting germ cell fate. Nat. Rev. Mol. Cell Biol. 16: 406–416.

Tropepe V., S. Hitoshi, C. Sirard, T. W. Mak, J. Rossant et al., 2001 Direct neural fate specification from embryonic stem cells: a primitive mammalian neural stem cell stage acquired through a default mechanism. Neuron 30: 65–78.

Tursun B., T. Patel, P. Kratsios, and O. Hobert, 2011 Direct conversion of *C. elegans* germ cells into specific neuron types. Science 331: 304–308.

Updike D. and S. Strome, 2010 P granule assembly and function in *Caenorhabditis elegans* germ cells. J. Androl. 31: 53–60.

Updike D. L., S. J. Hachey, J. Kreher, and S. Strome, 2011 P granules extend the nuclear pore complex environment in the *C. elegans* germ line. J. Cell Biol. 192: 939–948.

Updike D. L., A. K. Knutson, T. A. Egelhofer, A. C. Campbell, and S. Strome, 2014 Germ-granule components prevent somatic development in the *C. elegans* germline. Curr. Biol. 24: 970–975.

Voronina E., G. Seydoux, P. Sassone-Corsi, and I. Nagamori, 2011 RNA granules in germ cells. Cold Spring Harb. Perspect. Biol. 3: a002774.

Wang X., Y. Zhao, K. Wong, P. Ehlers, Y. Kohara et al., 2009 Identification of genes expressed in the hermaphrodite germ line of *C. elegans* using SAGE. BMC Genomics 10: 213.

Yan D., Z. Wu, A. D. Chisholm, and Y. Jin, 2009 The DLK-1 kinase promotes mRNA stability and local translation in *C. elegans* synapses and axon regeneration. Cell 138: 1005–1018.

